# High-fat diet-induced obesity increases susceptibility to endogenous and ethyl carbamate-induced somatic mutagenesis

**DOI:** 10.1101/2023.06.14.543760

**Authors:** Rebecca Lichtler, Mark J. Wilson, Jeffrey K. Wickliffe

## Abstract

Obesity and dietary fat consumption are associated with increased risk for certain cancers. We conducted a set of experiments designed to investigate both the inherent mutagenicity of high-fat diet (HFD) and the effect of HFD exposure on sensitivity to ethyl carbamate, a known mutagen. Greater body mass in females was associated with higher frequency of Pig-a mutant erythrocytes and HFD-induced obese female mice exhibited increased sensitivity to ethyl carbamate-induced mutagenesis compared to control mice. Together, these findings indicate that HFD exposure induces both endogenously generated changes related to mutagenesis and interacts with the mutagenic effects of ethyl carbamate.

## 1. Introduction

Obesity and overweight are highly prevalent in the United States and are associated with a number of conditions, including certain cancers. Of all new cancer cases diagnosed in 2015, 55% of those diagnosed in women and 24% of those diagnosed in men can be linked to obesity[1]. In fact, incidence of obesity-related cancers rose 7% between 2005 and 2014, while non-obesity related cancers declined by 13%, and the fastest growing obesity-related cancer type during that period was liver cancer, which rose by 29% [1,2]. Women, in particular, appear to be at increasing risk for obesity related cancers. Among women, rates of obesity-related cancers rose between 2001 and 2018, while cancers due to other risk factors declined or remained steady during the same time period [3]. Women with higher body mass index (BMI) were less likely to survive liver cancer following treatment than those with BMI considered to be within the normal range [4]. Along with evidence that higher BMI or adiposity increases cancer risk, dietary composition appears to play a role as well. High dietary fat consumption was associated with a 50% increase in the likelihood of having breast cancer in women [5]. Among men and women, higher consumption of dietary fat was associated with a 60% increase in colorectal cancer[6]. A meta-analysis of prospective cohort studies suggest consuming a diet high in fat, and saturated fat in particular, was associated with a 7-14% increase in lung cancer risk [7]. Another meta-analysis of prospective cohort studies that tracked liver cancer incidence found a 34% increased risk of liver cancer among those with the highest saturated fat consumption [8]. A recent study of patients with clonal hematopoiesis of indeterminate potential (CHIP), a phenomenon linked to hematologic cancers where blood cells accumulate mutations, uncovered obesity as risk factor for this condition[9].

The pathway from a genotoxic exposure, either dietary or chemical, to cancer development involves several stages, the first of which is initiation, or the establishment of a somatic gene mutation [10]. Despite the abundance of population-based association studies that link dietary fat consumption and cancer risk, very few controlled experimental studies have been conducted to evaluate the mutagenicity of HFD alone. Additionally, there is a dearth of studies that investigate the effects of co-exposure to HFD and environmental mutagens[11]. We have previously shown that high-fat diet (HFD) exposure in male C57BL/6J mice leads to higher frequency of spontaneous somatic mutations using the Pig-a gene mutation assay [12]. The Pig-a gene mutation assay was developed to measure the frequency of somatic gene mutations in peripheral red blood cells, by exploiting the glycosylphosphatidylinositol (GPI) cell surface anchor protein, found ubiquitously on red blood cells[13–16]. GPI anchor biosynthesis is dependent on a functional *Pig-a* gene, located on the X chromosome, meaning a single mutation can result in a GPI-deficient cell phenotype, and the frequency of GPI-deficient red blood cells can be enumerated using flow cytometric analysis[17].

Here we present the results from a set of experiments investigating the mutagenicity of HFD-induced obesity including both male and female mice for the first time. We hypothesized that HFD-induced obesity will increase the risk for both endogenous, spontaneous somatic mutagenesis and exogenous mutagen-induced mutagenesis. To test this hypothesis, we conducted two experiments, one designed to determine the effect of HFD alone (60% calories from fat) over time in both males and females on frequency of spontaneous somatic mutations using the Pig-a gene mutation assay. We also designed a two-factorial experiment whereby mice who had undergone long-term HFD exposure were challenged with oral ethyl carbamate and assessed for Pig-a gene mutation frequency after a sufficient mutation manifestation period. Ethyl carbamate is a known mutagen and carcinogen that is produced in food and beverage fermentation and has been shown to induce somatic gene mutations using the Pig-a gene mutation assay [16,18]. While the general public is not at high risk, those that regularly consume stone fruit brandy or fermented soy products may have a lifetime tumor risk of 0.01%, 100-times the risk that is generally considered acceptable [19–21]. Ethyl carbamate is converted into more toxic epoxide-containing compounds by esterases and the cytochrome P450 superfamily of phase I metabolic enzymes, several members of which are known to be perturbed in the context of obesity [22]. Together, this set of experiments explores both the endogenous and exogenous risk of somatic mutagenesis, one of the key components of the pathway to carcinogenicity, in the context of diet-induced obesity.

## 2. Methods

### 2.1 General animal care and analysis

Mice were reared at the Tulane University Downtown Vivarium on a 12-hour light/dark cycle. Research staff supplied C57BL/6J mice with HFD containing 60% calories from fat from Research Diets, Inc. (D12492) or with a low-fat diet (LFD) containing 10% calories from fat (D12450J), *ad libitum*. The LFD formulation serves as the most appropriate control to the HFD formulation because it is source-matched and contains equivalent sucrose content. Water and veterinary care were provided by vivarium staff. Unless otherwise stated, Prism software (GraphPad; San Diego, CA, USA) was used to conduct statistical analysis and to generate figures. Unless otherwise indicated, data in figures is presented as the arithmetic mean ± standard error with individual data points. Statistical significance was determined using a threshold of α=0.05, and results of statistical mean comparison tests are presented as the t statistic with degrees of freedom in subscript, followed by the corresponding p value.

### 2.2 Biochemical assays

To ensure that long-term HFD exposure had generated a physiologically obese phenotype in our model, plasma leptin, alanine amino transferase (ALT), and aspartate aminotransferase (AST), and total liver triglycerides were quantified using commercially available kits within the vehicle-control (deionized water) treated mice from the ethyl carbamate challenge experiment. Enzymatic activity of AST and ALT was measured as the synthesis of glutamate or pyruvate, respectively, that occurred per minute of incubation with each substrate at 37°C by Aspartate Aminotransferase Activity Assay Kit (Abcam ab105135) and the Alanine Transaminase Activity Assay Kit (Abcam ab105134). Plasma leptin was measured with the Mouse/Rat Leptin Quantikine ELISA Kit (R&D Systems, MOB00). Liver triglycerides were measured with the Triglyceride Quantification Assay kit (Abcam, ab65336), following tissue lysis and isolation of lipids with homogenization and boiling in a solution of 5% NP-40 in deionized water. All colorimetric and fluorescent results were analyzed using a Tecan Infinite 200 Pro and i-control software (V1.9, Mannedorf, Switzerland).

### 2.3 HFD-only experiment

#### 2.3.1 Animal care and sample collection

This experiment was conducted under the approval of Tulane University IACUC protocol 546. Mice were maintained on HFD or LFD conditions from 8 weeks old until 52 weeks old. Each dietary group consisted of 12 males and 12 females. Blood was taken at three time points, when mice were 17, 33 and 52 weeks of age. This sampling method was accomplished through repeated submandibular venipuncture under isoflurane anesthetization [23].

#### 2.3.2 Erythrocyte Pig-a mutant frequency analysis

Each blood sample was immediately transferred to a K_2_EDTA-coated microtainers (BD, 365974) and kept on ice. Samples were packed into Whirl-Pak® bags and shipped to Litron Laboratories (Rochester, NY). Pig-a mutant frequency analysis was carried out by Litron research staff following *In Vivo* MutaFlow® kit protocols [12,16,24].

#### 2.3.4 Statistical analysis

Body mass and mutant frequency were analyzed using mixed effects analysis of variance (ANOVA) for repeated measures with Bonferroni’s post hoc multiple comparisons test performed at each time point between LFD and HFD subjects. Simple linear regression analysis of the relationship between body mass and Pig-a mutant frequency was also conducted at each time point, and p value for slope deviation from zero and R^2^ correlation coefficient were reported.

### 2.4 Ethyl carbamate challenge experiment

#### 2.4.1 Animal care and sample collection

This experiment was conducted under approved Tulane University IACUC protocol 36-105. 48 males and 48 females were reared on HFD or LFD from 8 weeks old until they reached 42 weeks old, when the experiment was terminated. Nine weeks prior to termination, at 33 weeks, mice were divided into groups of 4-6, and exposed to 0, 100, 200, or 400 mg/kg/day of ethyl carbamate (CAS # 51-79-6; Sigma, U2500, Lot # WXBC3505V) in deionized water via oral gavage for 3 consecutive days following the procedure outlined by Labash et al. [16]. At 42 weeks, to maximize yield and quality of blood samples, mice were weighed and then anesthetized with no more than 5% isoflurane (v/v) and 100% oxygen saturation at a 1 L/min flow rate. The sternal cavity was opened for full visualization of the heart, and blood was collected from the left ventricle with a heparin-coated 22G needle and 1 mL syringe. The blood was then transferred to K_2_EDTA-coated microtainers (BD, 365974) to prevent coagulation. Immediately after collection, 60 μL of each sample was transferred to a tube containing an anticoagulant solution provided by Litron Laboratories and kept on ice for further processing following the Mouse MutaFlow^PLUS^ Kit manual (Litron Laboratories, v170925).

#### 2.4.2 Pig-a mutant frequency analysis

Pig-a mutant frequency analysis was conducted at Tulane University using a modified MutaFlow method. In summary, white blood cells and platelets were removed with Lympholyte®-Mammal (CedarLane, CL5110) and remaining reticulocytes, erythrocytes, and platelets were labeled with Anti-CD24-PE (Miltenyi, 130-110-688) and Anti-CD61-PE (Miltenyi, 130-102-628) antibodies and anti-PE magnetic microbeads (Miltenyi, 130-048-801). A small portion of each sample was retained for cell density determination and the remainder was enriched for mutant cells via immunomagnetic separation on LS columns (Miltenyi, 130-042-401) within a magnetic field. SYTO 13 nucleic acid dye (Invitrogen, S7575) was added to discriminate mature erythrocytes from immature reticulocytes and remaining lymphocytes. Total mature erythrocyte and Pig-a mutant erythrocyte counts were enumerated via the Miltenyi MACSQuant Analyzer 10 flow cytometer. Pig-a mutant mature erythrocytes were classified as cells exhibiting absence of the PE fluorescence signal and intracellular SYTO 13, indicating a lack of both CD24 surface markers and nucleic acids. Mutant frequency was calculated as the number of mutant mature erythrocytes per 10^6^ total mature erythrocytes.

#### 2.4.3 Statistical analysis

Pig-a mutant frequency was analyzed using two-way ANOVA with diet and ethyl carbamate dose as the two independent factors, with Bonferroni’s post hoc comparison test between LFD and HFD groups within each dose group. A dose-response curve was plotted to compare the effect of ethyl carbamate dose on Pig-a mutant frequency between LFD and HFD groups, with slope comparison and R^2^ calculation as well as an F test to compare dose-response curve slopes between HFD and LFD groups. Males and females were analyzed separately.

## 3. Results

### 3.1 Biochemical markers of obesity

HFD exposure in the untreated group of the ethyl carbamate challenge experiment (vehicle controls) led to elevated liver triglycerides in females (t_8_=2.409, p=0.0425) and males (t_8_=2.942, p=0.0186) and elevated plasma leptin in females (t_10_=7.187, p<0.0001) and males (t_10_=4.717, p=0.00082) (Fig 1A-B). Activity of liver enzymes AST (females: t_6_=2.857, p=0.0289; males: t_6_=3.841, p=0.0086) and ALT (females: t_6_=3.354, p=0.0153; males: t_6_=5.574, p=0.0014) were elevated in plasma, as well (Fig 1C-D). These biochemical findings indicate HFD exposure until 42 weeks of age was sufficient to generate increased body mass, fatty liver, and elevated circulating levels of liver enzymes and a circulating adipokine, all characteristic features of HFD-induced obesity.

**Fig 1.**
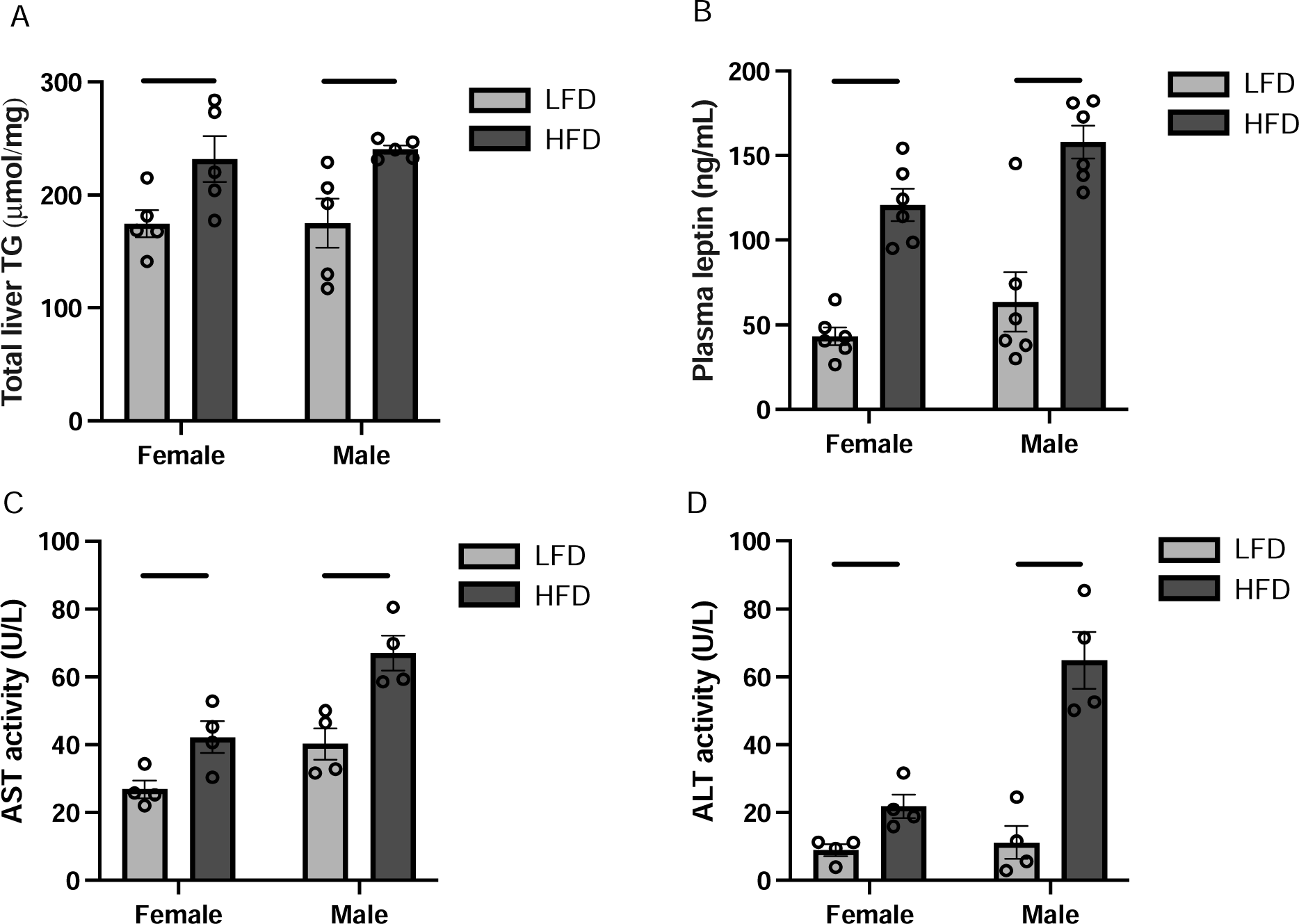
Effect of HFD-exposure on biochemical markers of obesity at 42 weeks-of-age. (A) Total liver triglycerides, (B) plasma leptin, (C) plasma AST activity, and (D) plasma ALT activity. * p<0.05, ** p<0.01, *** p<0.001, **** p<0.0001 per two-tailed Student’s t test between LFD and HFD groups, conducted within males and females separately (n=4-6).

### 3.2 HFD-only experiment

To assess the mutagenicity of long-term HFD exposure, the erythrocyte Pig-a mutation assay was conducted with sequential blood sampling at 17, 33 and 52 weeks old. By week 52, one LFD female was euthanized due to injury, and two HFD males died due to undetermined causes. Both HFD males and HFD females were significantly heavier than their LFD counterparts at each sampling timepoint. On average, HFD females were 6.98 (t_17.11_=6.334, p<0.0001), 19.99 (t_15.12_=8.281, p<0.0001), and 32.10 (t_14.92_=13.27, p<0.0001) grams heavier than LFD females at 17, 33, and 52 weeks, respectively (Fig 2A). On average, HFD males were 10.62 (t_14.24_=8.151, p<0.0001), 14.83 (t_11.34_=11.18, p<0.0001), and 15.90 (t_11.07_=9.494, p<0.0001) grams heavier than LFD males at 17, 33, and 52 weeks, respectively (Fig 2B).

**Fig 2.**
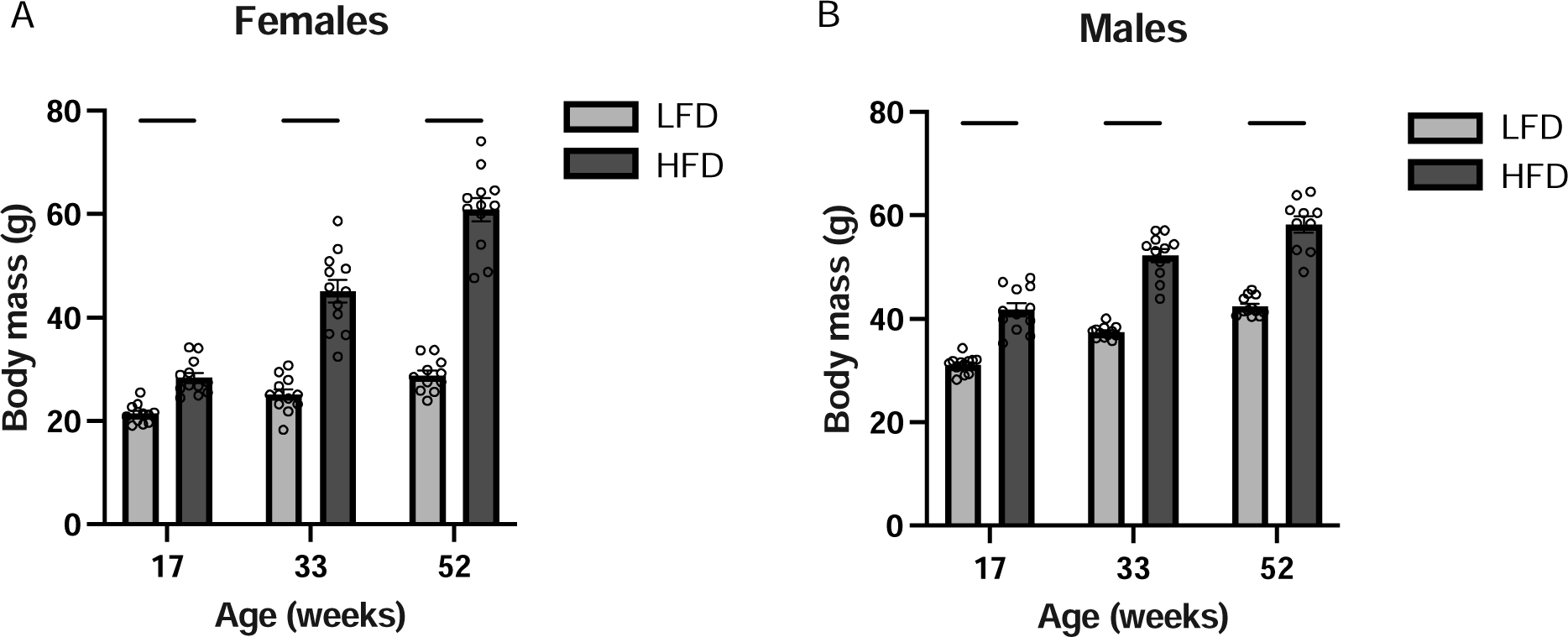
Effect of diet on body mass at each sampling timepoint. Male (A) and female (B) body mass in LFD and HFD groups at 17, 33 and 52 weeks of age. **** p<0.0001 per mixed effects analysis (for repeated measures) with Bonferroni’s multiple comparisons test, between LFD and HFD groups at each timepoint, conducted in males and females independently (n=10-12).

One extreme outlier LFD female was excluded from all Pig-a mutant frequency analyses due to very high frequency of Pig-a mutant erythrocytes detected at 33 and 52 weeks (>4 standard deviations). Blood samples that did not pass quality control parameters determined by the operators at Litron Laboratories were not reported. Both raw and fold-difference erythrocyte mutant frequency values were reported to normalize interexperiment variability. Despite a trend of an approximately 1.5-2-fold elevation, there was no significant difference in mutant frequency at any sampling timepoint between LFD and HFD females (Fig 3A-B). Because of the variability in body mass observed in response to HFD and the stochastic nature of somatic mutagenesis, simple linear regression was performed to determine a dose-response curve for the association between body mass and mutant frequency at each sampling time point. In females, simple linear regression for the association between body mass and erythrocyte mutant frequency per million total erythrocytes indicated the slope was not significantly different from zero at 17 weeks, indicating the was no association between body mass and mutation frequency at the first sampling timepoint (Fig 3C, Table 1). By 33 weeks, there was significant positive association between body mass and mutant frequency, with an R^2^ of 0.2004 and significantly non-zero slope (p=0.0322), indicating mice with a higher body mass were more likely to experience more frequent somatic mutations (Fig 3D, Table 1). By 52 weeks the association between body mass and mutant frequency remained positive, with an R^2^ of 0.2518 and a significant slope (p=0.0286) (Fig 3E, Table 1), indicating higher body mass was associated with higher incidence of somatic mutations, and this association grew stronger with age in females. Among males, while there was a similar trend towards higher (∼2-fold) mutant frequency among the HFD group, there was no significant differences between the LFD and HFD groups at any time point (Fig 4A-B), and there was no significant correlation between body mass and mutant frequency at any sampling timepoint (Fig 4C-E, Table 2).

**Figure 3.**
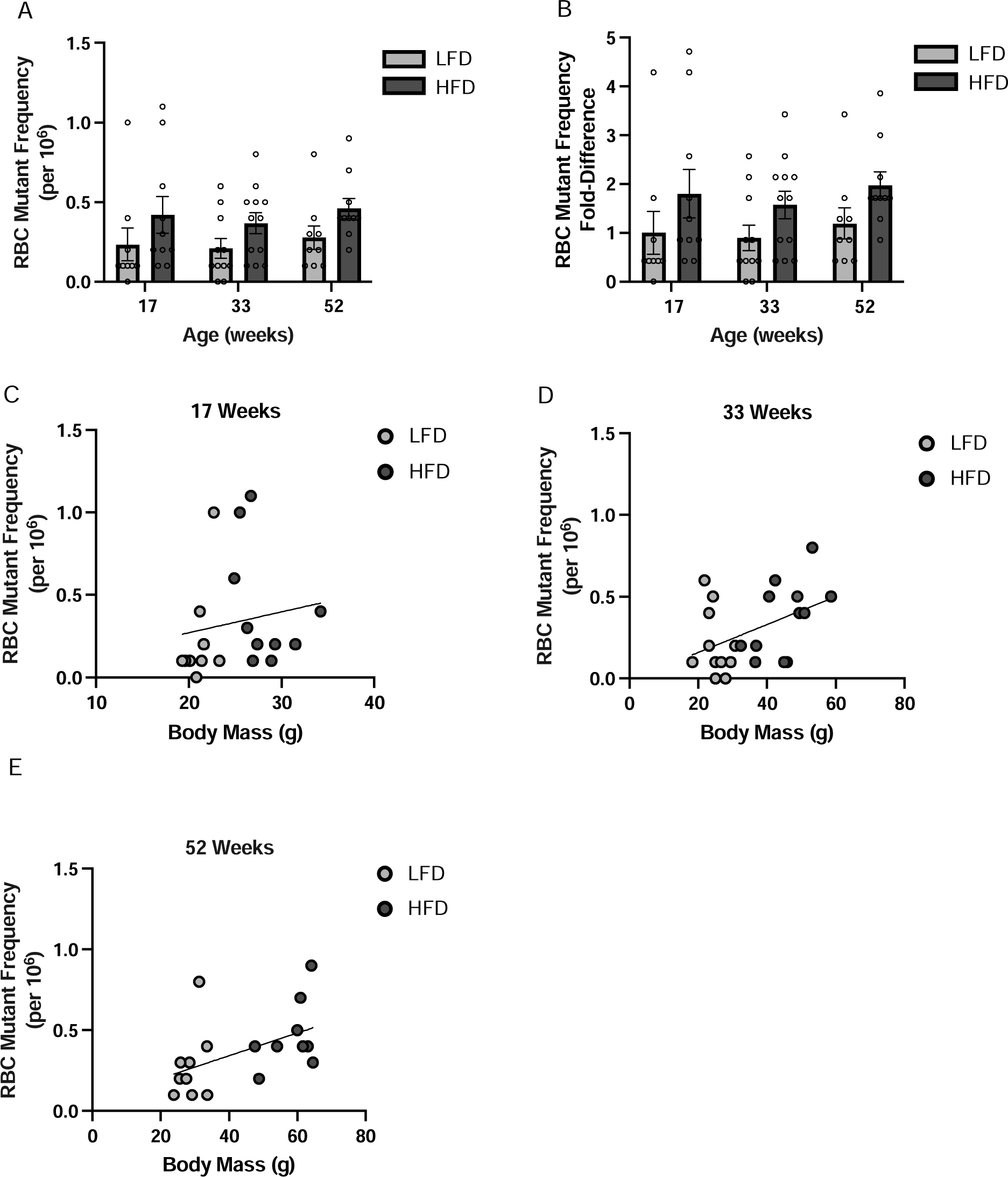
Effects of diet and body mass on spontaneous mutation frequency in erythrocytes at 17, 33 and 52 weeks of age in females. (A) Mutant frequency per million erythrocytes, and (B) Mutant frequency fold-difference between LFD and HFD groups at each time point, per mixed effects analysis (for repeated measures) with Bonferroni’s multiple comparisons test, between LFD and HFD group sat each timepoint (n=10-12). (C) Correlation between body mass and erythrocyte mutant frequency per million cells at 17 weeks. (D) Correlation between body mass and erythrocyte mutant frequency per million cells at 33 weeks. (E) Correlation between body mass and erythrocyte mutant frequency per million cells at 52 weeks.

**Fig 4.**
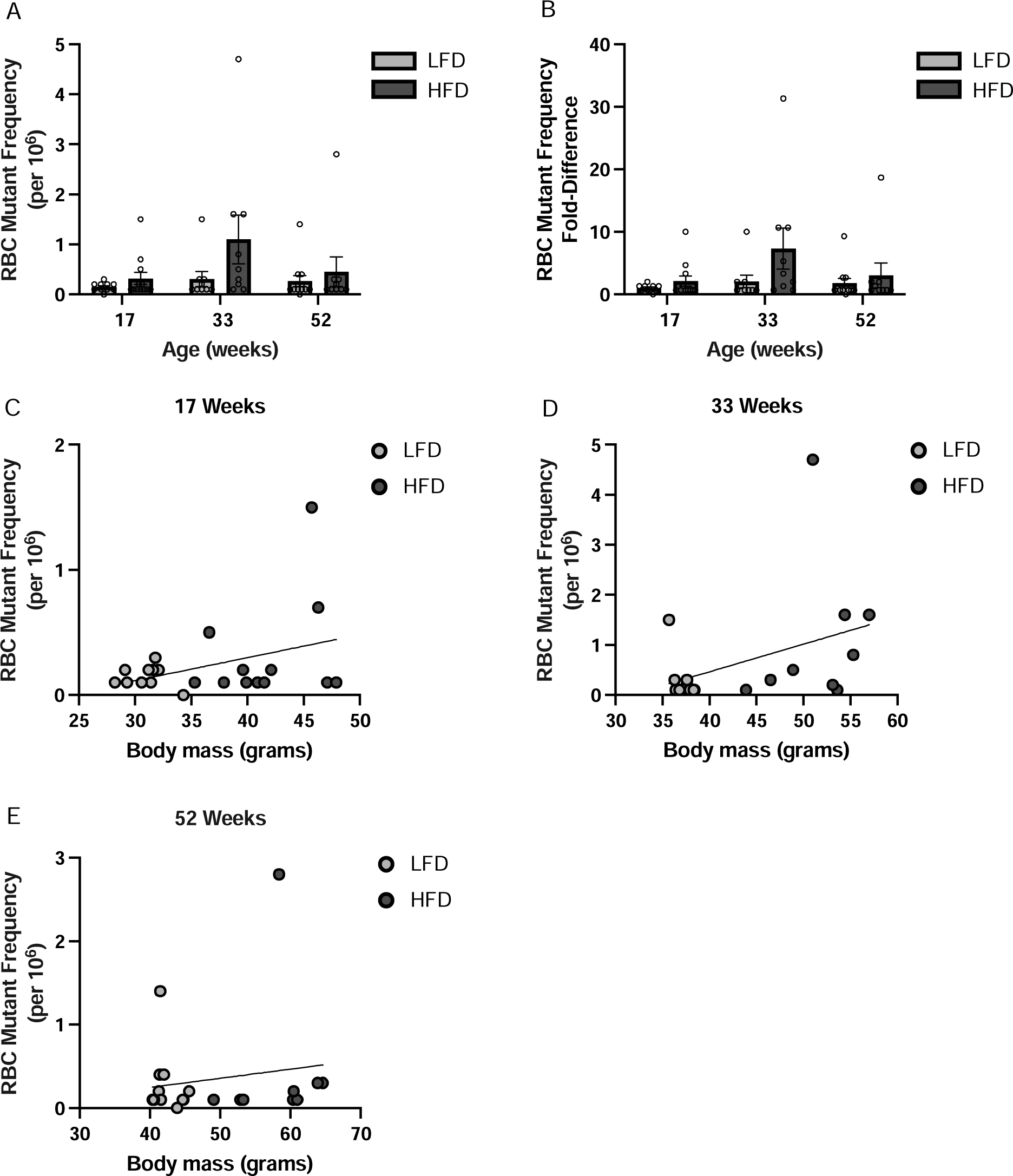
Effects of diet and body mass on spontaneous mutation frequency in erythrocytes at 17, 33 and 52 weeks of age in males. (A) Mutant frequency per million erythrocytes, and (B) Mutant frequency fold-difference between LFD and HFD groups at each time point, per mixed effects analysis (for repeated measures) with Bonferroni’s multiple comparisons test, between LFD and HFD group sat each timepoint (n=10-12). (C) Correlation between body mass and erythrocyte mutant frequency per million cells at 17 weeks in males. (D) Correlation between body mass and erythrocyte mutant frequency per million cells at 33 weeks in males. (E) Correlation between body mass and erythrocyte mutant frequency per million cells at 52 weeks in males.

**Table 1.**
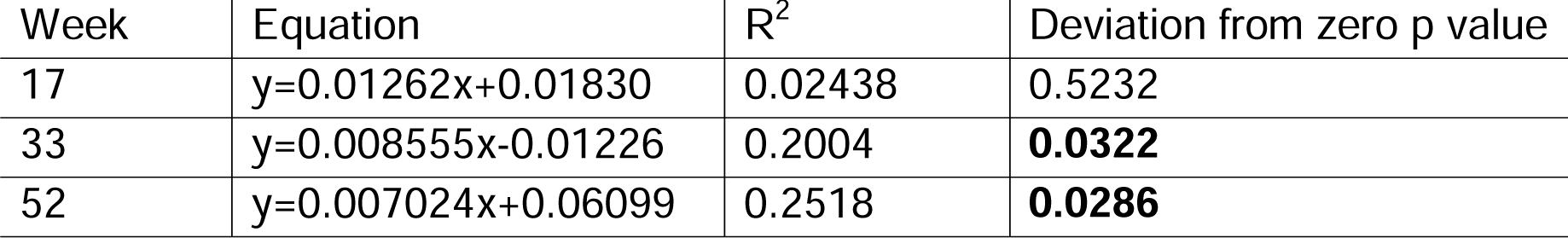
Simple linear regression correlating body mass and Pig-a mutant frequency in females after 17, 33 and 52 weeks of dietary exposure. Bold type indicates the slope deviates significantly from zero (α=0.05).

**Table 2.**
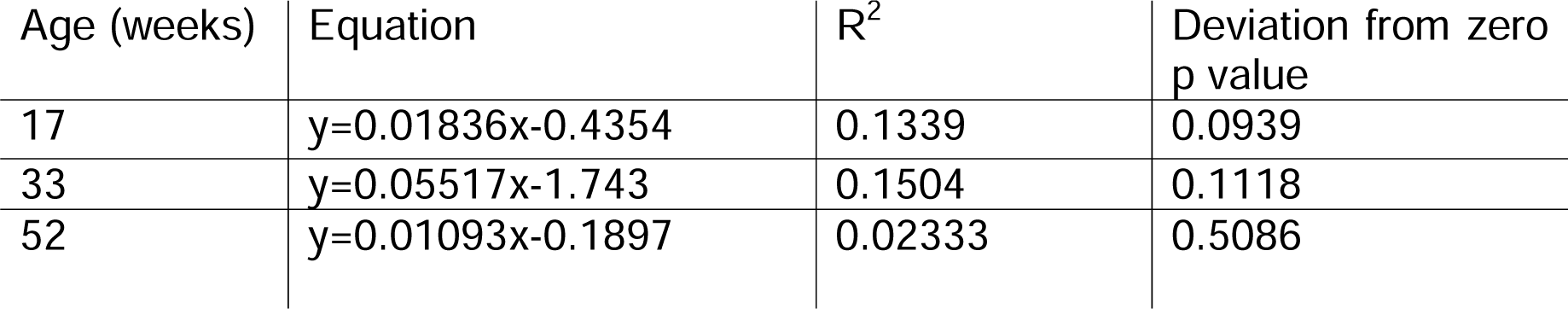
Simple linear regression correlating body mass and Pig-a mutant frequency in males after 17, 33 and 52 weeks of dietary exposure.

### 3.3 Ethyl carbamate challenge experiment

To determine the effect of HFD on sensitivity to a known mutagen, mice were reared on HFD or LFD starting at 7 weeks of age, exposed to ethyl carbamate at 33 weeks, and the experiment was terminated following a 9-week mutation manifestation period. Dietary conditions were maintained for the entirety of the experiment, and 93 of 96 total mice survived until termination at 42 weeks old. One female HFD mouse was sacrificed at 17 weeks due to excessive pruritus with unhealed dermal lesions and lack of weight gain. One male HFD mouse was sacrificed at 22 weeks due to sudden rapid weight loss, several large obstructive growths were observed in the colon during necropsy. One female LFD mouse was cannibalized at 29 weeks. All HFD groups were significantly heavier than their corresponding LFD controls and there was no significant effect of ethyl carbamate dose group on final body mass, (p<0.0001 for all LFD to HFD comparisons) (Fig 5).

**Fig 5.**
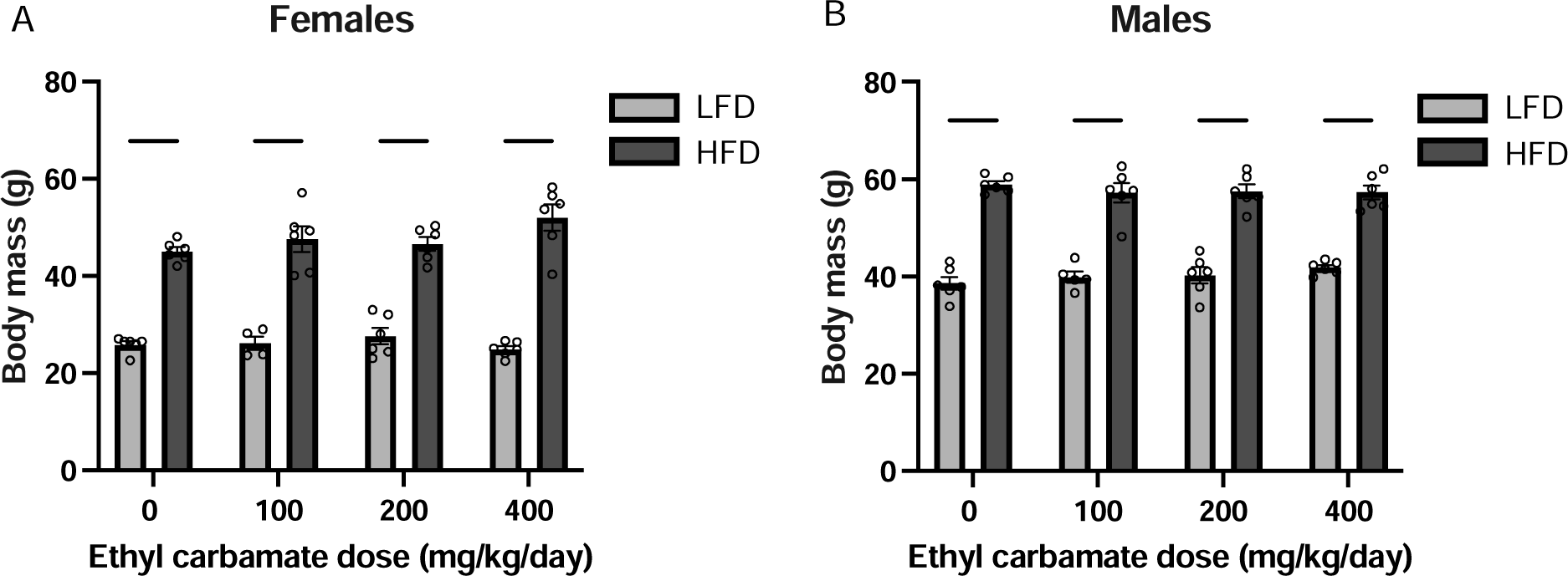
The final body mass among all dietary and ethyl carbamate treatment groups. (A) final body mass in males, and (B) final body mass in males. **** p <0.0001 per two-way ANOVA with Bonferroni-corrected comparison between all groups (only those comparisons within dose groups are shown here).

HFD exposure in females led to increased susceptibility to somatic mutations induced by exposure to 400 mg/kg/day ethyl carbamate (t_38_=4.67, p=0.0001; Fig 6A). The difference was on average 3.8-fold higher in the HFD group than the LFD group, within those who received 400 mg/kg/day ethyl carbamate (t_38_=4.67, p=0.0001; Fig 6B). To determine if HFD females were more sensitive than LFD females to the same ethyl carbamate dose regimen, dose response curves were plotted with regression analysis between ethyl carbamate dose and Pig-a mutant frequency response. Simple linear regression indicated the dose-response slope does not significantly differ from zero in the LFD group (p=0.3872) and there was no correlation between ethyl carbamate dose and mutant frequency response (R^2^=0.0376; Fig 6C, Table 3). The HFD regression curve is highly significantly different from zero (p<0.0001), and there is a strong correlation between ethyl carbamate dose and mutant frequency (R^2^ =0.5924; Fig 6C, Table 3). The slope of the HFD curve is significantly higher than the LFD curve, indicating HFD exposure significantly increases the susceptibility of female mice to ethyl carbamate induced somatic mutagenesis, and that the two dietary treatment groups display different dose-response curves in response to ethyl carbamate (p=0.0003; Table 3).

**Fig 6.**
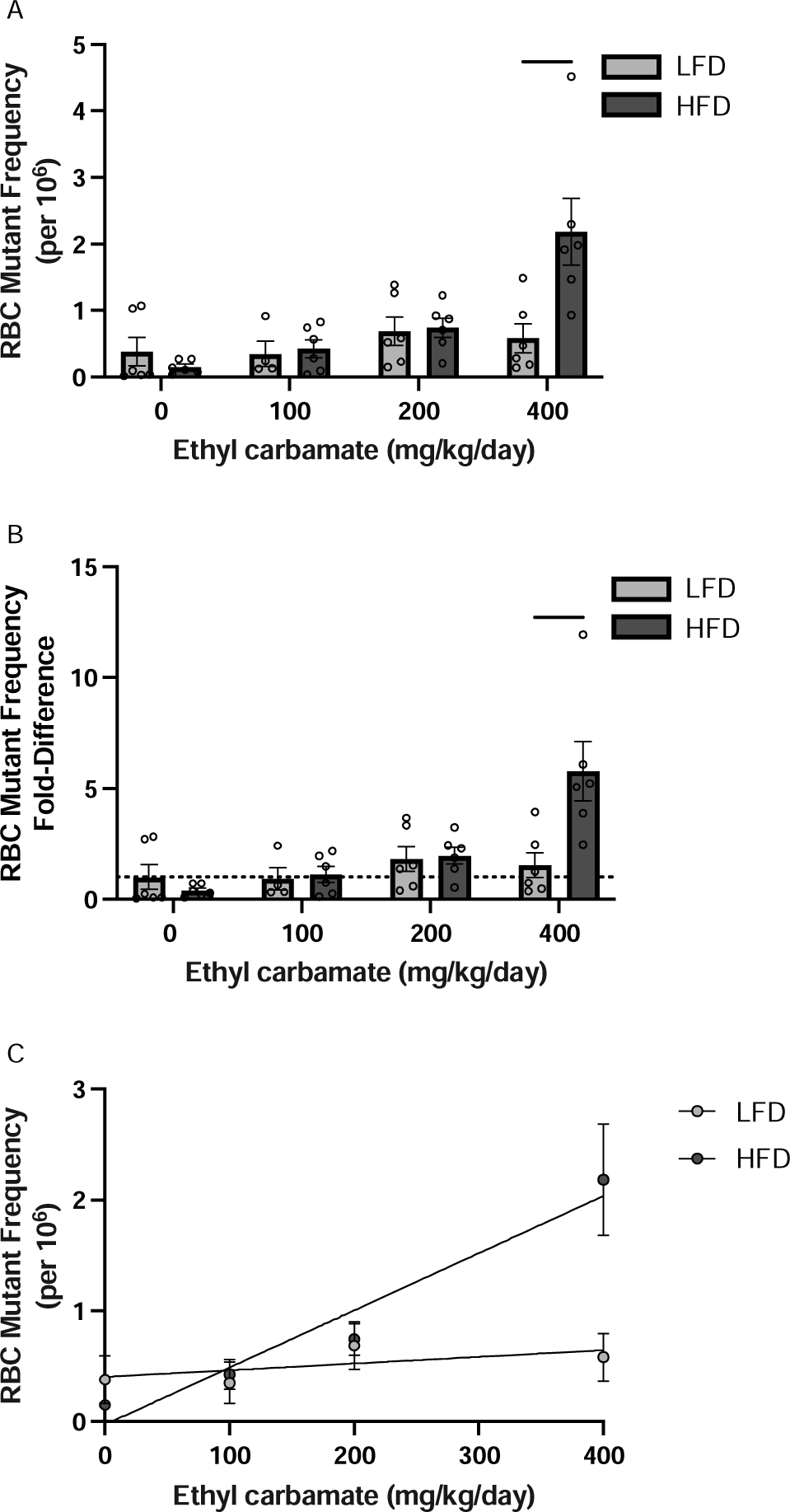
Effects of diet and body mass on mutant frequency in erythrocytes of female mice exposed to 0, 100, 200, and 400 mg/kg/day ethyl carbamate. (A) Mutant frequency per million erythrocytes in each ethyl carbamate dose group. (B) Mutant frequency fold-difference between LFD and HFD groups in each ethyl carbamate dose group. (C) Dose-response curves for ethyl carbamate dose and erythrocyte mutant frequency with simple linear regression analysis, points represent mean ±standard error. *** p<0.001 per two-way ANOVA with Bonferroni’s multiple comparison test between LFD and HFD groups (n=4-6).

**Table 3.** Simple linear regression dose-response analysis correlating ethyl carbamate dose and Pig-a mutant frequency response in females.

| Diet | Equation | $R^2$ | Deviation from zero<br>p value | Slope comparison<br>p value |
| --- | --- | --- | --- | --- |
| LFD | $y = 0.0006029x + 0.4032$ | 0.0376 | 0.3872 | **0.0003** |
| HFD | $y = 0.005166x - 0.02898$ | 0.5924 | **<0.0001** | |

Bold type indicates the slope or slope difference deviates significantly from zero ( $\alpha=0.05$ ).

Among males there was no significant difference between the response of the HFD and LFD groups to any level of ethyl carbamate treatment (Fig 7A). Though there was a trend suggesting HFD males exhibit an approximately 2-fold increase in Pig-a mutant frequency, regardless of ethyl carbamate treatment, these differences were not statistically significant (Fig 7B). Linear regression analysis of the correlation between ethyl carbamate dose and mutant frequency with slope comparison between the HFD and LFD groups suggest a positive trend, although the slopes were not significant and the R^2^ values were low (LFD: p=0.1032, R^2^=0.1486; HFD: p=0.0951, R^2^=0.1471; Fig 7C). There was ultimately no difference between the slopes for the LFD and HFD curves, suggesting both dietary groups are equally sensitive to ethyl carbamate-induced mutagenesis (p=0.5654, Table 4).

**Fig 7.**
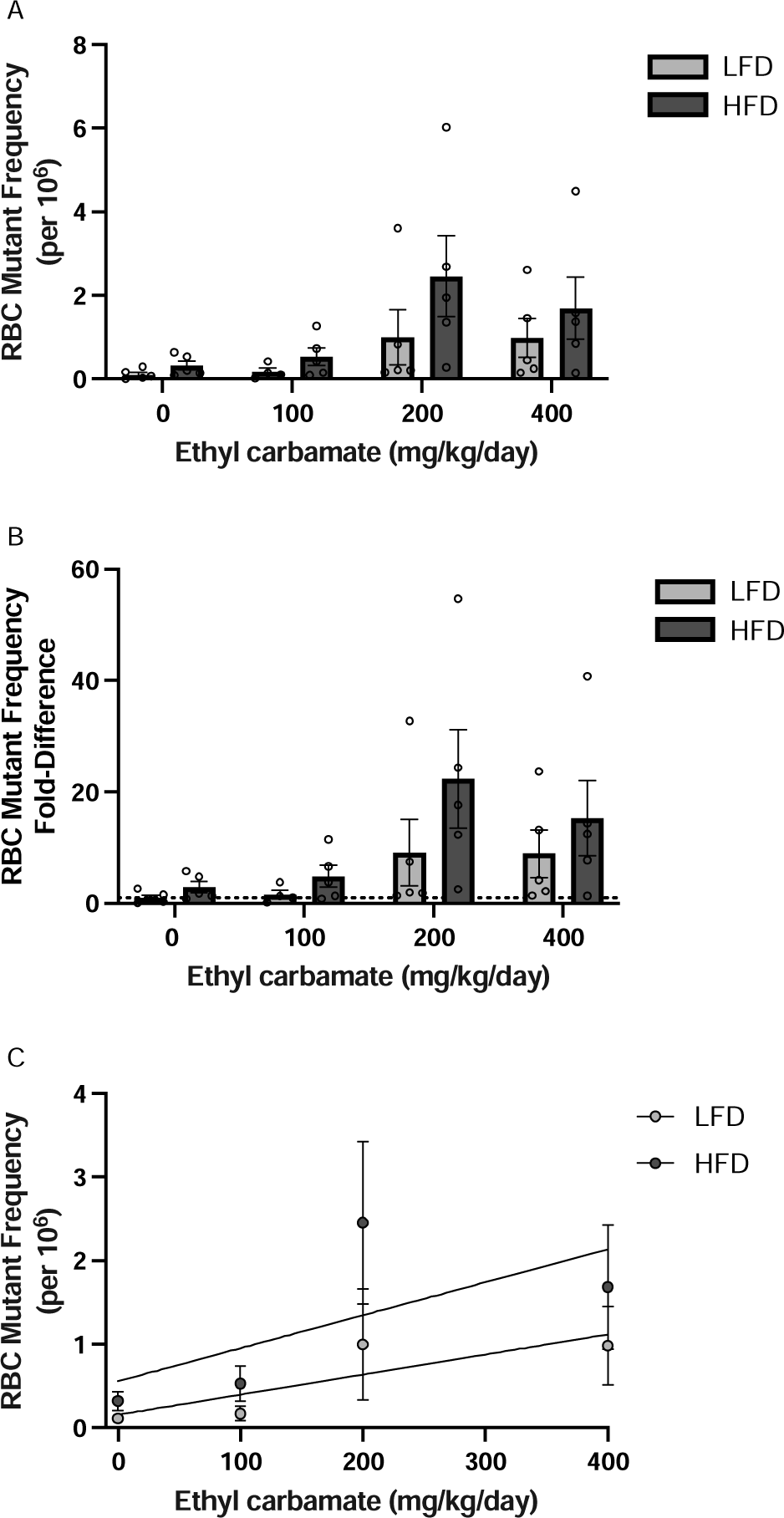
Effects of diet and body mass on mutant frequency in erythrocytes of male mice exposed to 0, 100, 200, and 400 mg/kg/day ethyl carbamate. (A) Mutant frequency per million erythrocytes in each ethyl carbamate dose group. (B) Mutant frequency fold-difference between LFD and HFD groups in each ethyl carbamate dose group. (C) Dose-response curves for ethyl carbamate and erythrocyte mutant frequency with simple linear regression analysis, points represent mean ±standard error (n=4-5).

**Table 4:** Simple linear regression dose response analysis between ethyl.

| Diet | Equation | R<sup>2</sup> | Deviation from zero<br>p value | Slope comparison<br>p value |
| --- | --- | --- | --- | --- |
| LFD | $y=0.002403x+0.1553$ | 0.1486 | 0.1032 | 0.5654 |
| HFD | $y=0.003947x+0.5569$ | 0.1471 | 0.0951 | |

**carbamate dose and Pig-a mutant frequency response in males.**

## 4. Discussion

The purpose of this set of experiments was to evaluate the mutagenicity of HFD both alone and with a chemical mutagen challenge using the Pig-a gene mutation assay. The findings presented here describe the first time that the effect of HFD on spontaneous somatic gene mutation frequency was tested in both males and females with repeated sampling over the lifespan. Interestingly, females in this long-term experiment were more responsive to HFD-induced weight gain as they aged than males, as indicated by the fact that HFD females were twice as heavy as LFD females, while HFD males were less than 50% heavier than LFD males after they reached 1 year of age (Fig 2). This was likely due to the fact that even males on the LFD become overweight over time while females do not, a phenomenon well-documented in the C57BL/6J strain[25]. Because of this tendency of males to gain weight over time, regardless of diet, we postulate that both LFD and HFD males represent some degree of biological overweight/obesity.

In females, linear regression analysis demonstrates that there is a significant dose-responsive relationship between body mass and Pig-a mutant frequency at 33 and 52 weeks old. This association between body mass and Pig-a mutant frequency strengthens with age and continued HFD exposure (Fig 3C-E, Table 1), suggesting that the length of the duration of HFD exposure over the lifespan is linked to degree of spontaneous gene mutation risk. These findings suggest that HFD-induced weight gain in females is sufficient to increase the risk of the initiation phase of carcinogenesis.

HFD females developed, on average, 3.8-times more somatic mutations in response to high-dose ethyl carbamate than LFD females (Fig 6B). In fact, obesity appears to potentiate the mutagenicity of ethyl carbamate in females, as indicated by the significance of the ethyl carbamate-mutant frequency dose-response slope among HFD, but not LFD, females (Fig 6C, Table 3). Among HFD males, mutagenicity of ethyl carbamate appears to saturate between 100 and 400 mg/kg/day, with a peak at 200 mg/kg/day (Fig 7A-C). Our previous findings suggested that HFD males exhibited a 2-fold elevation in mutation frequency, and that trend was recapitulated in this experiment, though the difference did not reach the threshold of statistical significance[12]. HFD males tended to exhibit non-significantly higher mutant frequencies than LFD males in each ethyl carbamate dose group, however the dose-response curves for LFD and HFD males shared a common slope (Fig 7C, Table 4). These results suggests that males on both diets were similarly sensitive to ethyl carbamate. We attribute the overall lack of association between diet and mutant frequency with the underlying weight gain observed in LFD males, and this supports our assertion that studies that only include male C57Bl6/J mice are not sufficient to explore the interactive effects of high-fat diet-induced obesity and environmental toxicants.

Further investigation is needed to elucidate the mechanism behind the increased ethyl carbamate sensitivity observed in HFD-exposed female mice. Several human studies have indicated that individuals with a higher body mass index metabolize xenobiotics differently. For example, the cytochrome P450 3A4 isoform tends to be less active in the context of obesity, while the 2e1 isoform tends to be more active. This group of xenobiotic metabolizing enzymes, part of the phase I class of chemical metabolism, can be responsible for both detoxifying and increasing the toxicity of drugs and other chemicals, depending on the substrate and efficiency of downstream phase II metabolizers. Our findings emphasize the importance of investigating the mechanisms underlying the HFD-induced mutagen sensitivity phenotype.

Interpretation of sex differences observed in this set of experiments is complicated by the different trends in weight gain between male and female mice. That is, males on the LFD gained weight consistently throughout the duration of the study, while females on the LFD maintain proportionally lower body mass. Increased weight gain by LFD males maintained for longer periods of time may mask differences that might be evident a study that included in truly lean (non-overweight, non-obese) males as a control group. Additional calorie restriction or reduced dietary fat may be required to generate a male equivalent of the female non-obese phenotype within this model system.

## 5. Conclusions

The findings presented here emphasize the importance of studying obesity as a mutagen and carcinogen and the necessity of including both sexes in diet-induced obesity research. For chemical risk assessment to properly protect the most vulnerable groups, obesity must be factored in. In our female model, obesity appears to act to both directly induce *de novo* mutations and to increase sensitivity to xenobiotic-induced mutagenesis. We demonstrated that obesity is likely involved in the initiation (mutagenesis) stage of cancer progression, however, further studies are needed to investigate the differences in male and female response to HFD exposure. The underlying the mechanism behind the association between HFD and ethyl carbamate exposure and somatic mutagenesis is also important to investigate further.

## Acknowledgements

We thank Stephen Dertinger, Dorothea Torous, and Svetlana Avlasevich of Litron Laboratories for conducting the Pig-a mutation assay for the long term HFD experiment. All funding for these projects was from intramural sources at Tulane University.

